# Development of a stable transformation method for *Saprolegnia parasitica*

**DOI:** 10.1101/2025.11.02.686010

**Authors:** Franziska Trusch, Israt Jahan Tumpa, Debbie McLaggan, Mark van der Giezen, Pieter van West

## Abstract

Saprolegniosis in salmonids, a disease caused by the oomycete *Saprolegnia parasitica*, poses a serious global threat to wild salmon and to aquaculture. To be able to functionally characterise genes in *S. parasitica*, it is essential to develop a stable transformation method for *S. parasitica*. We describe for the first time a method that can generate stable transgenic *S. parasitica* strains. Transformants were generated following the uptake and integration of a mutated gene from *Achlya hypogyna* conferring imidazole resistance, CYP51 using *S. parasitica* protoplasts in the presence of polyethylene glycol (PEG) and lipofectamine. This leads to production of CYP51 protein which catalyses a crucial demethylation step in the biosynthesis of ergosterol. As a result, there is no disruption of ergosterol synthesis and the transformants, but not the wild type *S. parasitica*, can grow in the presence of imidazole. Putative transformants growing in the presence of up to 10 mM imidazole were confirmed by PCR.

## 1. Introduction

Oomycetes are filamentous, spore-forming microorganisms that cause serious diseases in plants, fish, amphibians, and crustaceans, posing major threats to agriculture, aquaculture, biodiversity, and global food security (van West, 2006; Ghimire et al., 2021). Among them, *Saprolegnia parasitica* is a significant fish pathogen associated with severe losses in both wild and farmed salmonids worldwide (van den Berg et al., 2013; Minor et al., 2014). Saprolegniosis, the disease it causes, is characterized by white, cotton-like mycelial growth on fish skin, gills, and fins, often leading to mortality through osmotic and respiratory failure (van West et al., 2006; Sarowar et al., 2013). Reports suggest that around 10% of hatched salmon in aquaculture are affected (van West, 2006; Phillips et al., 2008), with rising incidences recently noted in Scottish hatcheries.

Understanding the pathogenicity of *S. parasitica* is crucial for developing effective control measures. While transient RNAi-based gene silencing has been used to study gene function (Saravia et al., 2014), a stable transformation system would enable more advanced functional gene characterisation and localization studies. Although successful transformation systems exist for several plant-pathogenic oomycetes, a stable method for *S. parasitica* had not yet been established. Among the available techniques, polyethylene glycol (PEG)-mediated protoplast transformation is considered the most reliable, involving protoplast generation, DNA uptake, and regeneration (Ghimire et al., 2021). This method has been successfully applied in other oomycete species such as *Phytophthora infestans* (Judelson et al., 1991), *Saprolegnia monoica* (Mort-Bontemps and Fèvre, 1997), and *Pythium aphanidermatum* (McLeod et al., 2008), but not however, for achieving stable transformation in *S. parasitica*.

We report here a novel approach to obtain stable transformants of *S. parasitica*. This watermould is sensitive to imidazole, an antifungal drug which inhibits cytochrome P450 14α-demethylases (CYP51). These are essential enzymes crucial for ergosterol synthesis. In yeast a dominant CYP51 mutant was shown to confer resistance to imidazole (Doignon et al., 1993). In this study, overexpression of a homologous CYP51 gene from *Achlya hypogyna* (an oomycete from the Saprolegniales order) is used as selection marker for increasing imidazole tolerance in *S. parasitica* to generate transformants. We have shown for the first time that through using this approach we can obtain stable transformants of *S. parasitica*.

## 2. Materials and Methods

### 2.1. Culture conditions of *S. parasitica* isolates

*S. parastica* isolate BB.36.08 (Shreves et. al. 2024) was used for this study. This strain was isolated from farmed Atlantic salmon (*Salmo salar*) in Scotland. The strain was routinely grown on Potato Dextrose Agar (PDA) media containing 24 g/L potato dextrose (Sigma-Aldrich) and 15 g/L agar (Formedium) at 18 °C. For the preparation of protoplasts and genomic DNA extraction, cultures were grown at 24 °C in pea broth (van West et al., 2010): 125 g/L birds eye peas; double autoclaved and filtered through cheese cloth after first autoclaving.

### 2.2. Construction of plasmids for transformation

CYP51 from *Achlya hypogyna*, including 842 bp upstream and 862 bp downstream of the gene, was synthesised (Eurofins) and cloned by Gibson assembly into pGEM-T Easy vector (Promega) between *Apa*I and *Aat*II restriction sites. Subsequently, a CYP51 Y132H mutant, unable to bind imidazole (Sagatova et al., 2016), was created by site directed mutagenesis (Q5 Site-Directed Mutagenesis Kit, NEB). Primers are listed in Table 1. Transformations were carried out using α-select chemically competent *E. coli* cells (BIOLINE, BIO-85029,) grown in Luria-broth (LB medium) at 37 °C overnight. Plasmid DNA purification for transformation was done with QIAGEN Plasmid Maxi Kit (cat. no. 12181, Germany), following the manufacturer’s instructions. Constructs were verified by Sanger sequencing.

**Table 1.**
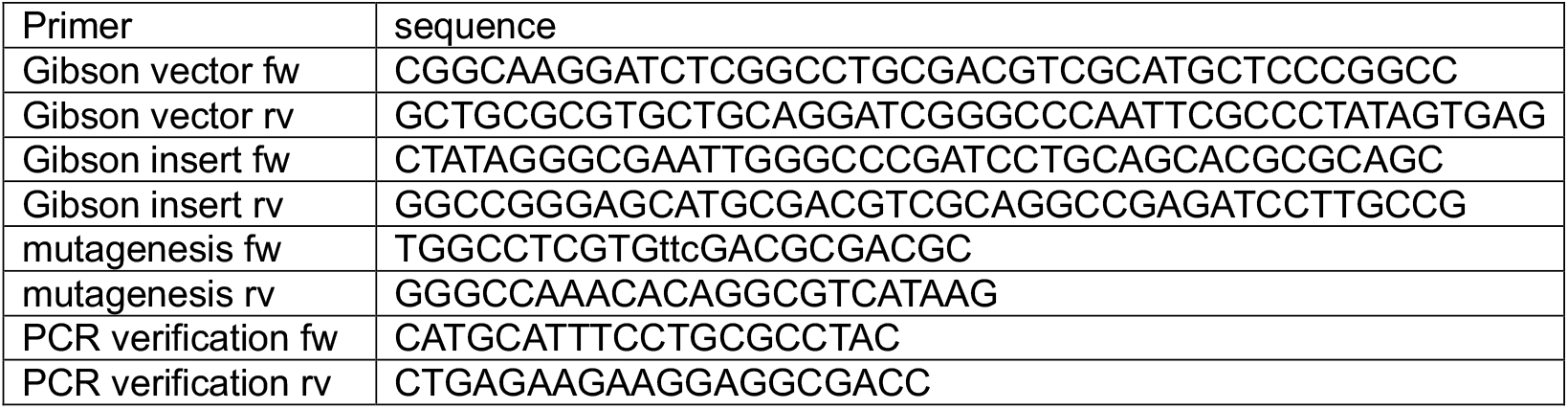
Primers used in this study.

### 2.3. Transformation procedure for *S. parasitica*

Development of PEG-mediated stable transformation system of *S. parasitica* was based essentially on the protocol by van West et al. (1998) for *P. infestans* and modified for *S. parasitica* as described below.

#### 2.3.1. Growth of *S. parasitica*

*S. parasitica* was grown on PDA plates (90 mm) for 4 to 5 days at 18 °C. The fluffy mycelium mat was then isolated from the plates (90 mm) containing PDA medium with a scalpel and chopped up and dispensed by pouring 40-60 mL pea broth into the plate (140 mm). The plate was incubated for 24 h at 18 °C. The young mycelial mats were then separated using a 70 µm mesh cell filter and the retained mycelia were transferred to 20 mL MES buffer (0.5 M Sorbitol, 20 mM KCl, 20 mM MES (pH 5.7), 10 mM CaCl_2_) in a falcon tube and washed twice. The mycelia were collected by filtration using a 70 µm mesh filter.

#### 2.3.2. Preparation of protoplasts for *S. parasitica*

To generate protoplasts, the harvested mycelial mats were treated with 20 mL MES buffer supplemented with 0.2 g cellulase from *Trichoderma viride* (Onuzuka, SERVA, Japan), and 0.1 g lysing enzymes from *T. harzianum* (Sigma). To maximise the protoplasts yield, the solution was incubated at room temperature with an incubation period of 30 mins with gentle shaking (55-60 rpm). The digested mycelial products were then strained through a mesh filter (70 µm) into a clean falcon tube to remove all debris. The filtrate was then passed through a smaller mesh filter (40 µm), followed by another 20 mL MES buffer to collect all the protoplasts in the filtrate. After centrifugation (4 min at 700 ×g) of the filtrate, the pellet of protoplasts was gently washed at room temperature in the following 3 buffers, with a centrifugation step in between (4 min at 700 ×g): 1) 10 mL MES buffer; 2) 10 mL 50:50 MES buffer/Sorbitol Tris (ST) solution (1 M sorbitol, 10 mM Tris, pH 7.5, 10 mM CaCl_2_); 3) 10 mL ST solution. Finally, the centrifuged protoplasts were resuspended in 970 µL ST solution and the protoplast concentration was determined. The concentration was typically about 1-3x10^7^/mL.

#### 2.3.3. Preparation of DNA/lipofectamine mix

The DNA/lipofectamine mixture was prepared by mixing of 85.7 µl Lipofectamine 3000 (Invitrogen, L3000-015) with 57.1 µl DNA containing 50-60 µg of CYP51 DNA for plasmid transformations.

#### 2.3.4. Transformation process

The protoplast suspension (around 1 mL) was combined with DNA/lipofectamine mix (142.8 µL) in a universal tube and then the tube was gently rotated at an angle for 30 sec. The tube was left for 5 min at room temperature then the transformation process was initiated by adding 1 mL PEG solution (PEG 3350 with 0.5 M Tris pH 7.5 and 100 mM CaCl_2_) on the side of the tube at the top of the protoplast suspension at an angle while rotating again for 30 sec then leaving for 2 min room temperature. After gently inverting once, the mixture was left for another 5 min then 2 mL of sugar pea broth (pea broth containing 1 M Sorbitol) was added to the mixture, left for 2 min then gently inverted once, then 6 mL sugar pea broth was added, and the tube was left for 5 min. This entire process was carried out at room temperature. Finally, the protoplast mixture (∼10 ml) was poured into a 60 mm plate containing 12 mL of sugar pea broth and the plate was incubated at 18 °C for 24 hours to allow the protoplasts to regenerate.

### 2.4. Propagation of transformants

The percentage regenerated protoplasts was determined to be about 5%. The regenerated protoplasts were collected by centrifugation (10 min, 1000 ×g) and resuspended in 6-7 mL sugar pea broth. An aliquot of the suspension (1 mL) was then very gently spread on PDA plates containing 7 mM imidazole as selective media and incubated at 18 °C for 7-9 days. The small colonies that appeared on 7 mM imidazole plates were then transferred to 10 mM imidazole PDA for final selection to avoid false-positive transformants. Overall, a transformation efficiency of 1 putative transformant per 1.5 µg plasmid DNA was achieved.

### 2.5. Screening the transformants

The putative transformants that grew on selective agar media were cultured in liquid pea broth at 24 °C for 48h. DNA from freshly grown mycelia was then isolated following the protocol described by Shreves et al. (2024) which was based on the phenol/chloroform protocol by Zelaya-Molina et al. (2011). In brief, fresh mycelia were ground by mechanical force with sterile beads into a Fastprep24 5G (MP Biomedicals™, Santa Ana, CA, USA) under liquid nitrogen, mixed with 800 µl of an extraction buffer (10 mM tris-HCI (pH 8), 50 mM EDTA, 0.5% (w/v) SDS, 0.5% (v/v) Triton X-100, and 0.5% (v/v) Tween 20) and 2 µl of 20 mg/mL proteinase-K (Promega, USA) and incubated for 30 mins at 55 °C, then cooled to room temperature. The aqueous phase was extracted by adding 800 µl phenol-chloroform-isoamyl alcohol (25:24:1) (ACROS Organics™, Belgium), mixing and then centrifugating (10 min, 10,000 ×g). The upper phase (1 mL) was removed and mixed with equal volume of molecular grade isopropanol (Fisher Scientific, USA) and placed at -20 °C for 15 mins. The DNA was then precipitated by centrifugation at 10,000 ×g for 10 min at 4 °C. The DNA pellet was washed with molecular grade 70 % (v/v), centrifuged (1 min, 10,000 ×g) then washed again with 100 % ethanol (Sigma Aldrich), then finally collected by centrifugation (1 min, 10,000 ×g), air dried and resuspended with DNase/RNase-free water. The concentration and purity of isolated DNA was determined by nanodrop and 1% agarose gel electrophoresis. Part of the CYP51 gene was amplified by PCR using the genomic DNA isolated from the transformants (see Table 1). This was performed by employing Go-Taq G2 polymerase (Promega). PCR conditions: initial heating (95 °C, 2 min); 34 cycles of denaturation (95 °C, 10 s); annealing (55 °C, 30 s); and extension (72 °C, 2 min) with a final extension for 10 min at 72 °C. DNA from the non-transformed wild type (WT) was used as a negative control and plasmid CYP51 as a positive control. The PCR product (471bp) was visualized using 1% agarose gel electrophoresis.

## 3. Results

### 3.1. Optimization of imidazole concentration for selection of transformants

To determine the lowest imidazole concentration resulting in growth inhibition a series of imidazole concentrations from 0 mM to 20 mM were tested for growth of *S. parasitica*. No growth was observed in the presence of 7 mM imidazole. A mock transformation was performed without DNA and protoplasts were cultured on PDA plates containing 0 mM to 7 mM imidazole. The mock ‘transformants’ were found to grow best from 0 mM to 4 mM imidazole (after 5 d), with growth declining significantly at and above 5 mM imidazole (**Figure 1**). At 7 mM imidazole mock transformants did not grow at all, or occasional growth was observed after 14 days when presumably the activity of imidazole had been reduced due to degradation in the agar. Following the mock transformation, it was decided to use 7 mM imidazole to select for transformants then subsequently subculture putative transformants in 10 mM imidazole containing media.

**Fig. 1.**
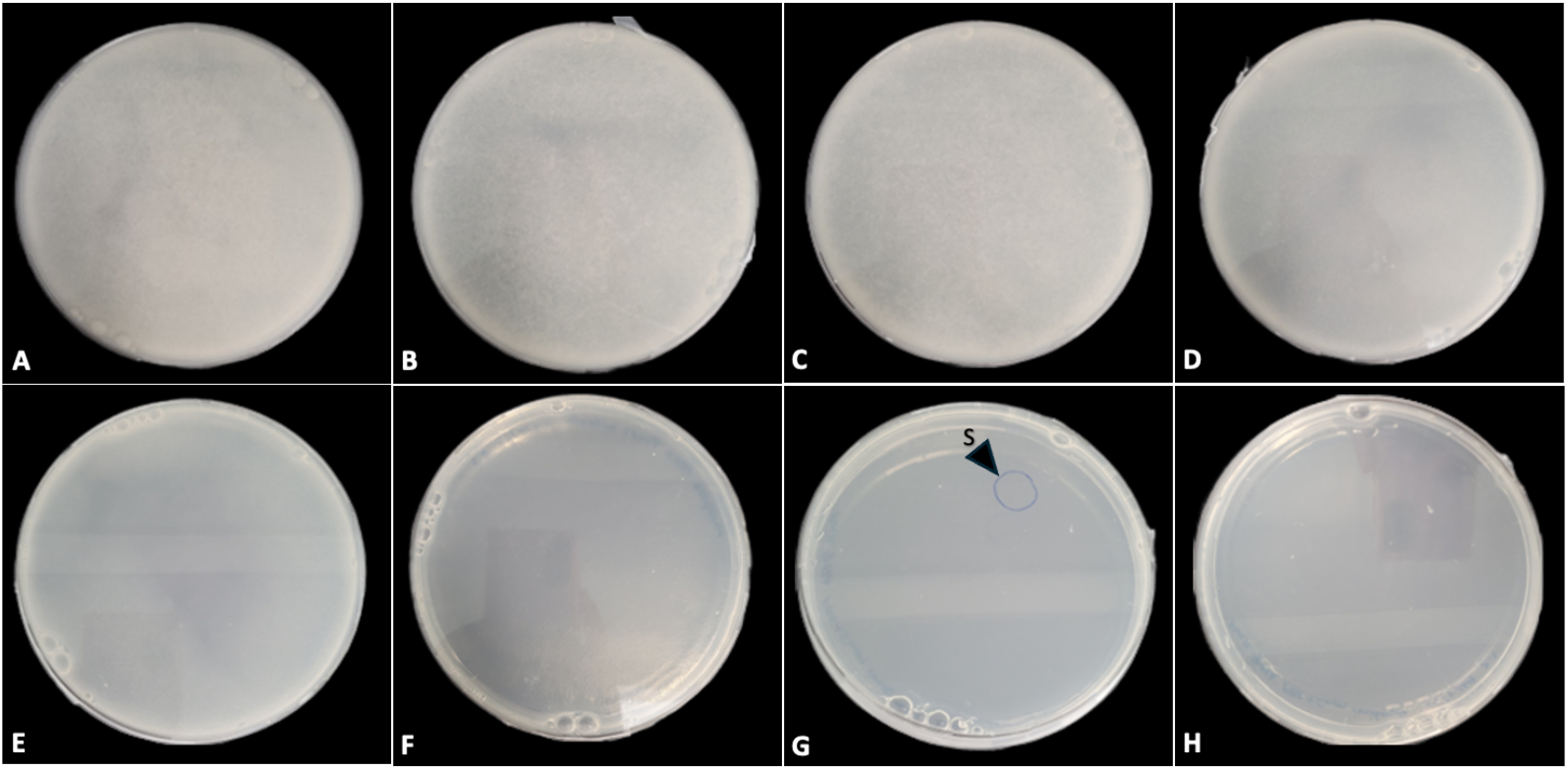
Testing series of concentration (0 mM to 7 mM) of Imidazole on regenerative protoplasts obtained from mock transformation of *S. parasitica* isolate (OQ678462). After 5 d of incubation, the panel shows: thick mycelial mat growing on PDA plates with **A**) 0 mM, **B**) 1 mM, **C**) 2 mM, **D**) 3 mM and **E**) 4 mM concentration of Imidazole. A thin mat of mycelia and a single colony (**S**) of mycelia were observed at plates **F**) and **G**) with 5 mM and 6 mM Imidazole, respectively. No growth was found in plate **H**) with 7 mM Imidazole.

### 3.2. Transformation of *S. parasitica* with CYP51-1

*S. parasitica* isolate BB.36.08 was transformed with plasmid CYP51-1. Selection was performed using 7 mM imidazole, then putative transformants were sub-cultured three times at 10 mM imidazole. An average of 20 imidazole resistant putative transformants were obtained per 50-60 µg of CYP51 DNA. Putative transformants were analysed by PCR (**Figure 2**). A DNA band of about 4.71 kb was amplified in the transformants and the positive control CYP51 plasmid DNA. The putative transformants were sub-cultured three times on 10 mM imidazole (**Figure 3**) prior to verifying stable incorporation of CYP51 DNA plasmid via PCR.

**Fig. 2.**
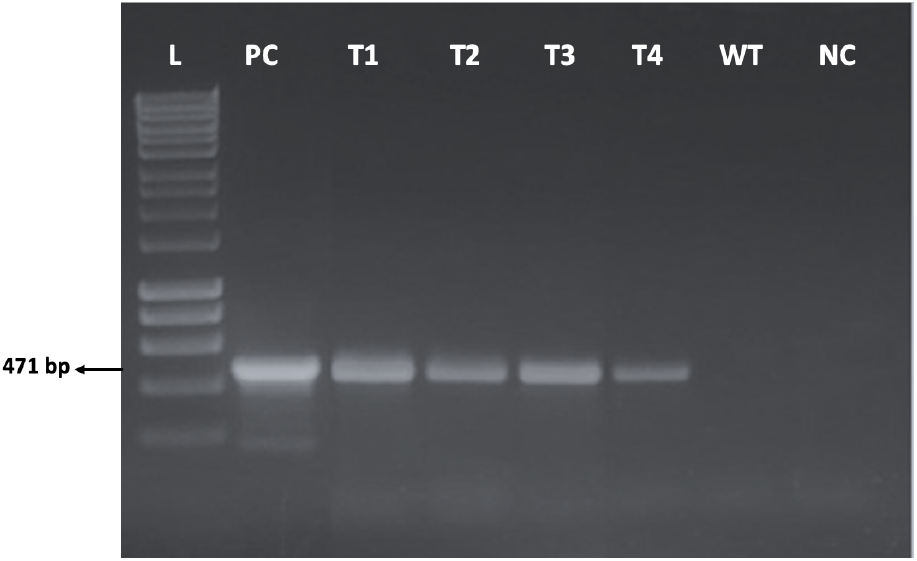
Qualitative PCR detection of CYP51 DNA isolated from putative transformants of *S. parasitica*. The panel represents DNA from CYP51-1 (**PC**) plasmid as positive control and DNA obtained from wild type (**WT**) *S. parasitica* isolate, and water only (**NC**) as negative controls. **T1, T2, T3**, and **T4** represents successful transformants with amplification of a 471 bp DNA similar as the positive control. **L** represents 1kb HyperLadder™ as molecular weight marker.

**Fig. 3.**
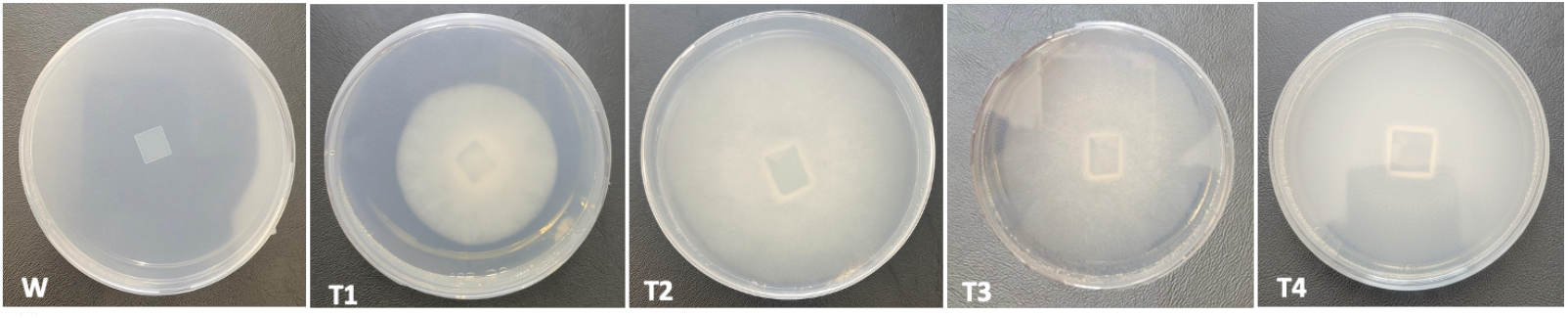
Selection of transformants with 7 mM Imidazole selection marker. **WT** refers wild type (no growth on selection plate) as negative control, and **T1, T2, T3** and **T4** represents successful transformants with visible mycelial growth with 10 mM imidazole PDA to avoid false-positive transformants.

## 4. Discussion

We describe the first effective PEG mediated transformation protocol for an oomycete animal pathogen, *S. parasitica* using a drug-resistant gene construct coupled with a reporter gene. In this study, one construct contained the coding sequences of an essential ergosterol synthesizing protein (CYP51). Verification of the integrated transgene was performed using PCR reactions.

Initially we tested several antibiotics and chemicals to determine if these would suppress growth in *S. parasitica*. These included: chloramphenicol, hygromycin, geneticin, gentamycin, puromycin, WR99210, trimethoprim, but none of these were suitable as selection markers. We then decided to investigate whether CYP51 would be suitable in combination with imidazole (**Figure 1**).

Our transformation method is based on generating protoplasts employing cellulase and lysing enzymes. A 30 min long lysis step using lysing enzymes from *T. viride* and *T. harzianum* facilitated in protoplasting with a good number of round and healthy-looking protoplasts with about 10% of the cells containing nuclei. The CYP51 protein was used as our target selection marker protein which plays a crucial step in biosynthesis of ergosterol - an important sterol for the cell membrane for Saprolegniales (Warrilow et al., 2014). *S. parasitica* is an oomycete which can produce CYP51 and can initiate the pathway of ergosterol synthesis (Warrilow et al, 2014), whereas this sterol pathway has been lost in most of the plant oomycetes, e.g. *Phytophthora* spp. (Madoui et al., 2009) The SpCYP51 protein in *S. parasitica* encodes a 14α-demethylase enzyme which binds to lanosterol (a precursor of ergosterol) and catalyses the demethylation step in conversion of lanosterol to ergosterol in the biosynthetic pathway (Warrilow et al., 2014). Antifungal drugs which are known to be effective in interfering with the competitive binding of 14α-demethylase enzyme to prevent conversion of lanosterol to ergosterol. This disruption in the biosynthetic process resulted into the accumulation of methylated sterol precursors (lanosterol) and depletion of ergosterol production and ultimately inhibits the cell growth and causes cell death (Barrs et al., 2024). According to Warrilow et al. (2014), imidazole-based azoles are 30% more effective to inhibit SpCYP51 *in vivo*, specially clotrimazole treatment in *S. parasitica* have been found to accumulate large amount (around 92% of the total sterols) of lanosterol in the cell membrane and resulted in depletion of ergosterol and other sterol synthesis. In this study, we have found that 7 mM imidazole is enough to inhibit cell growth of *S. parasitica*. During transformation, the insertion of a mutated form of the CYP51 gene from *A. hypogyna* leads to overexpression of this CYP51 dependent enzyme in the transformants. Therefore, there is minimal to no disruption of ergosterol synthesis, which results in these transformants override the resistance to 7mM concentration of imidazole as there is more CYP51 produced than in the non-transformed wild type. We decided to use the CYP51 gene from *A. hypogyna* rather from same *S. parasitica* to avoid the possibility of gene silencing of the endogenous and transformed CYP51 genes.

We obtained several transformants using imidazole selection system, but some of the transformants lost their incorporated vectors over time as has also been found in other transgenic oomycetes (Ghimire et al., 2010).

## 5. Conclusion

We have accomplished, for the first time, a method that can generate transgenic *S. parasitica* strains. The data presented in this paper should help with the development of stable transformation procedures for other *Saprolegnia* spp. and other animal oomycetes. This stable transformation protocol provides a basis for future research on the functional analysis of *S. parasitica* genes and the cell biology processes involved in the interaction between *S. parasitica* and its hosts.

## 6. Acknowledgement

The authors would like to show gratitude to the Commonwealth Scholarship Commission in the UK for funding this project, the BBSRC (APP19187) and the Norwegian Research Council (grant 301170). Furthermore, we would like to acknowledge Dr Marcia Saraiva, who has tested several antibiotics for possible transformation of *S. parasitica*.

## 8. CRediT authorship contribution statement

Franziska Trusch: conceptualisation, formal analysis, methodology, investigation, validation, writing, reviewing and editing. Israt Jahan Tumpa: formal analysis, methodology, investigation, data curation, validation, funding acquisition, writing. Debbie McLaggan: formal analysis, resources, project administration, writing, reviewing and editing. Mark van der Giezen: resources, funding acquisition, reviewing and editing. Pieter van West: supervision, conceptualization, formal analysis, methodology, resources, validation, project administration, funding acquisition, writing, reviewing and editing.

